# RTK signaling and epidermis remodeling instruct dendrite regrowth after pruning in the fly PNS

**DOI:** 10.64898/2026.10.11.758319

**Authors:** Neeraja Sanal, Ulrike Gigengack, Sebastian Rumpf

**Affiliations:** Multiscale Imaging Center and Institute for Neurobiology, University of Münster, Röntgenstrasse 16, 48149 Münster, Germany

**Keywords:** dendrite, pruning, regrowth, ecdysone, receptor tyrosine kinase, epidermis

## Abstract

Morphology and function of many neuron types must be adapted to the current developmental stage through neurite pruning and regrowth. Pruning and regrowth occur during defined temporal windows, but how these are defined is often unclear. Drosophila peripheral c4da neurons prune their larval sensory dendrites in an ecdysone-induced manner during metamorphosis. They subsequently regrow adult-specific dendrites, but how dendrite regeneration is linked to pruning is not known. We show that dendrite regrowth requires two receptor tyrosine kinases (RTKs), the insulin receptor InR and Ret, and present evidence that RTK ligands are derived from the adjacent epidermis. The Drosophila epidermis is extensively remodeled during metamorphosis through ecdysone-dependent apoptosis of larval epidermal cells and expansion of adult so-called histoblasts. We find that this occurs during a time window between dendrite pruning and regrowth, and that preventing epidermis remodeling inhibits dendrite remodeling. Furthermore, the two processes might also be evolutionarily linked. We propose that dendrite pruning and regrowth are part of a biphasic response to ecdysone, where a delayed regrowth signal is via the epidermis to ensure coordination of tissue developmental processes.

## Introduction

Developmental pruning, the regulated degeneration of neurites or synapses, is an important mechanism that serves to specify neuronal circuitry and to adapt neuronal circuits to different developmental stages (Riccomagno and Kolodkin, 2015, Rumpf et al., 2017, Rui, 2024). While neurite pruning often involves local microtubule disassembly (Rumpf et al., 2019) and mechanical forces acting on neurites (Krämer et al., 2023), synapse pruning requires actin disassembly (Frommeyer et al., 2026) and phagocytic synapse removal by glial cells (Scott-Hewitt et al., 2020). Developmental neuronal rewiring can then be achieved through regrowth of morphologically adapted neurites (Lee et al., 2000). Defects in developmental neuronal remodeling have been linked to neurological disorders like schizophrenia, underscoring its importance for normal brain development (Johnson and Stevens, 2018).

As developmental mechanisms, pruning and regrowth are under strict temporal control and occur in distinct phases, but how these are defined is often unclear. A good model to study pruning is the fruit fly Drosophila, where pruning occurs at large scale during metamorphosis. Here, the molting hormone ecdysone, a steroid hormone, triggers neurite pruning by inducing a specific transcriptional programme (Lee et al., 2000; Wlliams et al., 2005; Kuo et al., 2005). As Drosophila is a holometabolic organism, neuronal pruning serves to disassemble the larval nervous system and make space for new, adult-adapted neuronal connections that are subsequently generated through regrowth of axons and dendrites. How neurite regrowth is reinitiated after pruning is not generally established, but it has been suggested that it may depend on a transcriptional programme that leads to activation of the TOR growth pathway (Yaniv et al., 2012; Rabinovich et al., 2016).

The peripheral sensory class IV dendritic arborization neurons prune their long and branched larval dendrites within the first 16 hours after puparium formation (h APF) in an ecdysone-dependent manner (Williams et al., 2005; Kuo et al., 2005; Kirilly et al., 2009), and regrow them at a later stage that starts at around 40 h APF (Yoong et al., 2020; Sanal et al., 2023). The phosphoinositide 3-kinase (PI3K)/Target of Rapamycin (TOR) growth pathway plays an important role during both dendrite pruning and regrowth: during pruning, it must be inhibited in the neurons by a ubiquitin-dependent mechanism (Wong et al., 2013) and through counteracting kinases like the AMP-dependent protein kinase AMPK (Marzano et al., 2021; Yuh Chew et al., 2022), while it is required for dendrite regrowth (Sanal et al., 2023). During regrowth, TOR likely regulates the expression of cytoskeletal factors which are required for the process (Yoong et al., 2020; Sanal et al., 2023; Davies et al., 2025). The transcriptional pathway inducing axon regrowth in the fly central nervous system (Rabinovich et al., 2016) does not seem to be involved in dendrite regrowth in c4da neurons (Sanal et al., 2023), but whether other ecdysone-dependent pathways may be involved is not known. In addition, insulin signaling may be involved in dendrite regrowth, but the underlying logic is unclear (Sanal et al., 2023).

In addition to neurons, other larval tissues of Drosophila undergo remodeling during metamorphosis as well. Of note, the larval epidermis is composed of highly polyploid larval epidermal cells (LECs) to cope with the speed of larval growth. These cells undergo ecdysone-dependent apoptosis between 18 and 32 h APF and are replaced by diploid adult epidermal cells called histoblasts (Ninov et al., 2007). During this process, dying larval epidermal cells are extruded from the tight epidermal monolayer, while the histoblasts expand and take over the epidermis through cell divisions and migration until approximately 40 h APF (Michel and Dahmann, 2020). Interestingly, the extent of tissue remodeling varies between different taxa of holometabolic insects, and it is known that other dipterans such as mosquitos, which diverged from Drosophila ~ 250 million years ago (Arensburger et al., 2010), do not undergo extensive remodeling of their epidermis (Trager, 1937; Risler, 1959) or abdominal ganglia (Yin et al., 2024). Here, we investigated signaling cascades involved in c4da neuron dendrite regrowth. We find no evidence for cell autonomous ecdysone transcriptional signaling as a trigger. Instead, we find that two receptor tyrosine kinases, the insulin receptor and Ret, are required. Our data suggest that their respective ligands insulin-like peptide 6 (Ilp6) and Maverick (Mav) are derived locally, at least in part from the adjacent epidermis. Epidermis remodeling occurs during a time window between dendrite pruning and regrowth, and inhibition of epidermis remodeling strongly prohibits dendrite remodeling, demonstrating a tight link between the two processes. In further support of such a link, we find no evidence of pruning in sensory neurons of *Aedes* mosquitos which do not undergo epidermis remodeling. Thus, dendrite pruning and regrowth are coordinated with, and informed by, epidermis remodeling in Drosophila.

## Results

### Ecdysone is not required in a cell autonomous fashion for dendrite regrowth

C4da neurons prune their larval dendrites within the first 16 - 20 hours after puparium formation (h APF), and then regrow adult-specific dendrites starting at approximately 40 - 48 h APF (Sanal et al., 2023) (Fig. 1 A). As ecdysone is a major signal that induces morphological changes during metamorphosis and induces c4da neuron dendrite pruning, we wondered whether a second ecdysone signal could be involved in dendrite regrowth induction. To be able to manipulate ecdysone in a temporally-inducible fashion during the pupal stage, we made transgenes for the light-inducible shine-GAL4 system (di Pietro et al., 2021) under the c4da neuron-specific ppk promotor to generate *ppk-shine-GAL4*. As a functional test, we first asked if light-induced expression of dominant-negative ecdysone receptor (EcR DN) affects dendrite pruning. To this end, we grew larvae carrying UAS-EcR DN (or UAS-tdtomato as a control) in the dark until the late third instar stage. At approximately 24 hours before pupariation, we induced GAL4 activity with a short exposure to blue light, and then analyzed dendrite morphology at 20 h APF (Fig. 1 B). Dendrite pruning proceeded normally when animals carrying UAS-EcR DN were grown in complete darkness (Fig. 1 C, F), or when expression of UAS-tdtomato was induced before pupariation (Fig. 1 D, F). In contrast, strong pruning defects were observed when animals carrying UAS-EcR DN were exposed to blue light (Fig. 1 E, F), confirming that *ppk-shine-GAL4* is functional, and that ecdysone signaling is required for dendrite pruning.

**Figure 1.**
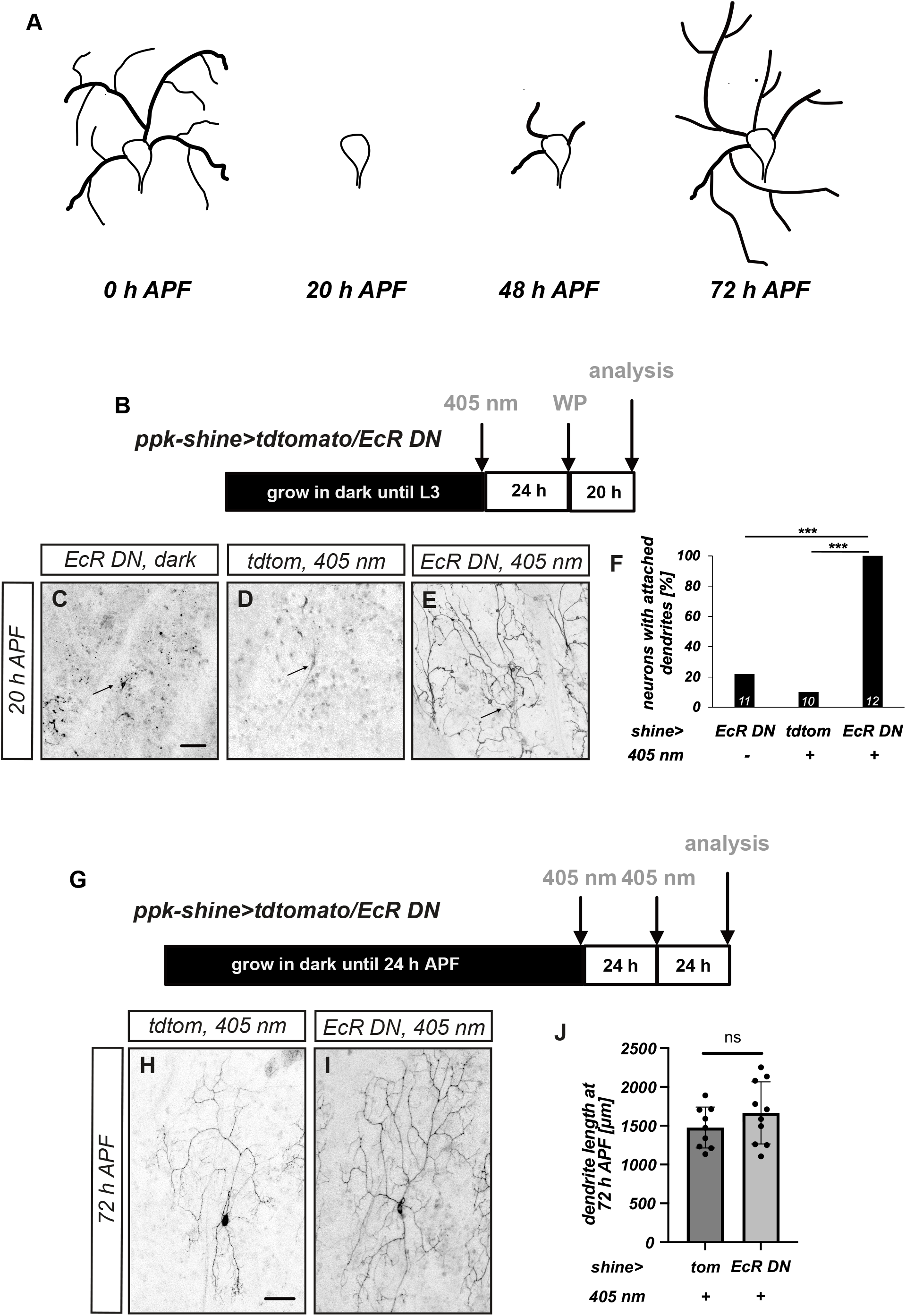
Cell-autonomous ecdysone signaling is required for c4da neuron dendrite pruning, but not for dendrite regrowth. **A** Schematic timecourse of c4da neuron dendrite remodeling. **B** – **F** Light-inducible ecdysone inhibition during c4da neuron dendrite pruning using *shine-GAL4*. **B** Timeline of experiments in **C** - **F**. Transgene expression was induced by blue light exposure 24 h before pupariation. Analysis of pruning phenotypes was at 20 h APF. C4da neurons were labeled with *ppk-CD4::tdGFP*. **C** Control c4da neuron in an animal carrying UAS-EcR DN without light exposure. **D** Control c4da neuron in an animal carrying UAS-tdtomato upon light exposure. **E** C4da neuron in an animal carrying UAS-EcR DN upon light exposure. **F** Penetrance of pruning defects in **C** - **E**. N=10-12 neurons, *** P<0.0005, two-tailed Fisher’s exact test. **G** - **J** Light-inducible ecdysone inhibition during c4da neuron dendrite regrowth. **G** Timeline of light-induced ecdysone inhibition in experiments. Transgene expression was induced by repeated blue light exposure at 24 h and 48 h APF, respectively. C4da neurons were labeled with *ppk-CD4::tdGFP*, and analysis of dendrite regrowth was at 72 h APF. **H** Control c4da neuron in an animal carrying UAS-tdtomato upon light exposure. **I** C4da neuron in an animal carrying UAS-EcR DN upon light exposure. **J** Quantification of total dendrite length in **H**, **I** at 72 h APF. N=9-10 neurons, n. s., not significant, Wilcoxon’s test. Scale bars in **C**, **H** are 50 µm.

To test whether ecdysone signaling is also required for dendrite regrowth, we next induced EcR DN or tdtomato as a control after the completion of pruning, but before dendrite regrowth. Here, we grew animals in the dark until after pupariation and exposed them to blue light at both 24 and 48 h APF for two minutes each time (Fig. 1 G). We then analyzed dendrite regrowth at 72 h APF and found that not only control animals expressing tdtomato (Fig. 1 H, J), but also the experimental animals expressing EcR DN (Fig. 1 I, J) had regrown long dendrites. Thus, cell autonomous ecdysone signaling is unlikely to be involved in the induction of dendrite regrowth after pruning.

### Ret is required for dendrite regrowth and interacts with InR/PI3K

We had previously shown that dendrite regrowth depends on PI3K/TOR signaling, and that knockdown of the insulin receptor InR caused (somewhat weaker) dendrite regrowth defects as well (Sanal et al., 2023). This suggested that RTKs might be upstream of TOR signaling, and potentially act as part of the regrowth-inducing mechanism. To test this idea further, we knocked down 10 candidate cell surface RTKs linked to neuron growth in c4da neurons with *ppk-GAL4* and asked if this affected dendrite regrowth (Fig. S1). This identified two dsRNA lines targeting Ret, a RTK required for morphogenesis of peripheral sensory neurons in mammals and flies (Luo et al., 2009; Hoyer et al., 2018) as well as gut maturation (Perea et al., 2017). Ret knockdown using two different dsRNA lines caused a significant reduction in dendrite length compared to controls (Fig. 2 A-D). Ret is expressed in larval c4da neurons (Hoyer et al., 2018). Using a GFP insertion in the Ret locus, we found that Ret was also continuously expressed in c4da neurons also between 0 - 72 h APF (Fig. S2). To independently confirm a role for Ret in dendrite regrowth, we used a transheterozygous *Ret* mutant and found that loss of Ret caused a similar degree of dendrite regrowth defects as the knockdown (Fig. 2 E, F, H). These dendrite regrowth defects of the *Ret* mutant could be rescued by re-expression of Ret under *ppk-GAL4* (Fig. 2 F-H).

**Figure 2.**
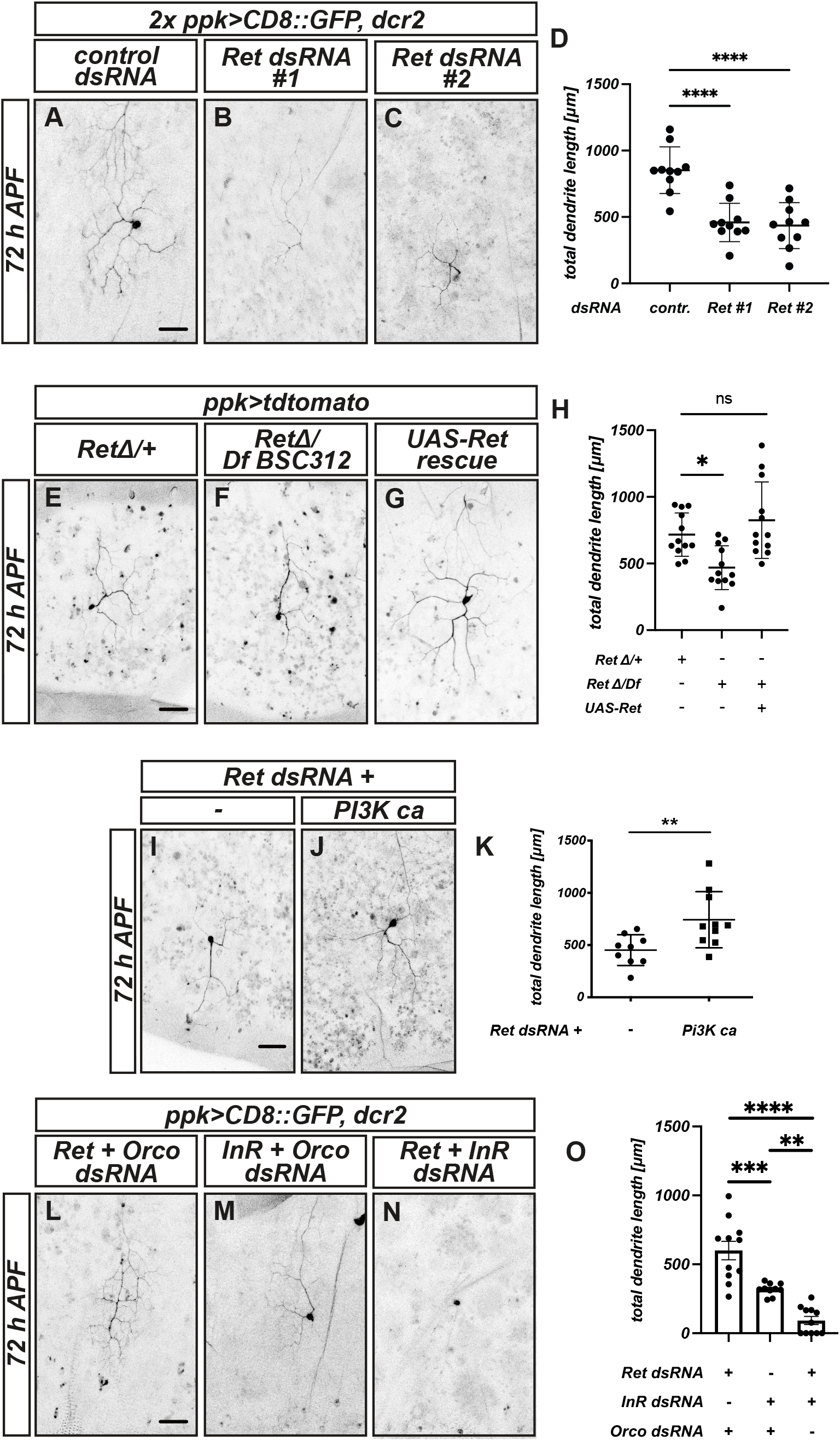
Ret is required for c4da neuron dendrite regrowth and interacts with PI3K/InR. **A** - **D** Ret knockdown causes dendrite regrowth defects. The indicated dsRNA constructs were expressed in c4da neurons under *ppk-GAL4*, and dendrite regrowth was assessed at 72 h APF. **A** C4da neuron expressing a control dsRNA directed against the Odorant co-receptor Orco. **B** C4da neuron expressing Ret dsRNA#1. directed against the Odorant co-receptor Orco. **C** C4da neuron expressing Ret dsRNA#2. **D** Quantification of total dendrite length in **A** – **C**. N= 10 neurons each, **** P<0.001, Wilcoxon’s test. **E** - **H** Dendrite regrowth defects in *Ret* mutant animals and cell-autonomous rescue. C4da neurons in animals of the indicated genotypes were labeled by tdtomato expressed under *ppk-GAL4*, and dendrite regrowth was assessed at 72 h APF. **E** C4da neuron in heterozygous *RetΔ*/+ animal. **F** C4da neuron in transheterozygous *RetΔ/Df BSC312* mutant animal. **G** C4da neuron in transheterozygous *RetΔ/Df BSC312* mutant animal carrying UAS-Ret. **H** Total dendrite length in **E** - **G**. N= 12 neurons each, n. s., not significant, * P<0.05, Wilcoxon’s test. **I** – **K** Constitutively active PI3K ameliorates dendrite regrowth defects caused by Ret knockdown. C4da neurons were labeled by CD8::GFP expression under *ppk-GAL4*, and dendrite regrowth was assessed at 72 h APF. **I** C4da neuron expressing Ret dsRNA#1. **J** C4da neuron coexpressing Ret dsRNA#1 and constitutively active PI3K-CAAX. **K** Total dendrite length in **I**, **J**. N=9-10, ** P<0.01, Wilcoxon’s test. **L** – **O** Simultaneous knockdown of Ret and InR causes strong dendrite regrowth defects. **L** C4da neuron coexpressing dsRNAs against Ret and Orco (as a control dsRNA). **M** C4da neuron coexpressing InR and Orco control dsRNAs. **N** C4da neuron coexpressing Ret and InR dsRNAs. **O** Quantification of total dendrite length in panels **L** - **N**. N=9-11 neurons, values are mean +/- s. d., * p<0.05, Wilcoxon’s test. Scale bars in **A**, **D**, **I** and **L** are 50 μm.

Ret can activate the PI3K pathway (Kim and Dressler, 2007), and we therefore speculated that it might work by activating this pathway during dendrite regrowth as well. Consistently, we observed that expression of PI3K^CAAX^, a membrane-targeted and therefore constitutively active form of PI3 kinase, caused a partial rescue of the regrowth defects caused by Ret knockdown (Fig. 2 I-K). As we had previously shown that Insulin Receptor knockdown also caused regrowth defects, we asked whether the two RTKs might act in a redundant manner to activate PI3K/TOR signaling during dendrite regrowth. Indeed, combined knockdown of Ret and InR caused very strong dendrite regrowth defects at 72 h APF (Fig. 2 L-O). We conclude that two RTKs, Ret and InR, play an important role in dendrite regrowth, likely by activating the PI3K/TOR signaling pathway.

### RTK ligands are linked to the epidermis

In order to better understand the activation of Ret and InR during dendrite regrowth, we next wanted to identify their relevant ligands. Assuming a non-cell autonomous pathway, we first knocked down candidate ligands linked to Ret or insulin signaling (Fig. S3) under the ubiquitously expressed *act5C-GAL4*, while concomitantly prohibiting GAL4 activity in c4da neurons with *ppk-GAL80*. Of 12 dsRNA lines targeting potential ligands, we identified one line targeting Drosophila Insulin-like peptide 6 (dIlp6) and two lines targeting Maverick (Mav), a TGFβ-like ligand for Ret (Hoyer et al., 2018, Myers et al., 2018), that caused partial c4da neuron dendrite regrowth defects at 72 h APF (Fig. 3 A-E). While the phenotypic strength of Mav knockdown roughly matched that of Ret knockdown, the effects of dILP6 knockdown seemed weaker than that of InR knockdown (compare Fig. 2). This could potentially be due to partial redundancy with other insulin-like peptides (see below).

**Figure 3.**
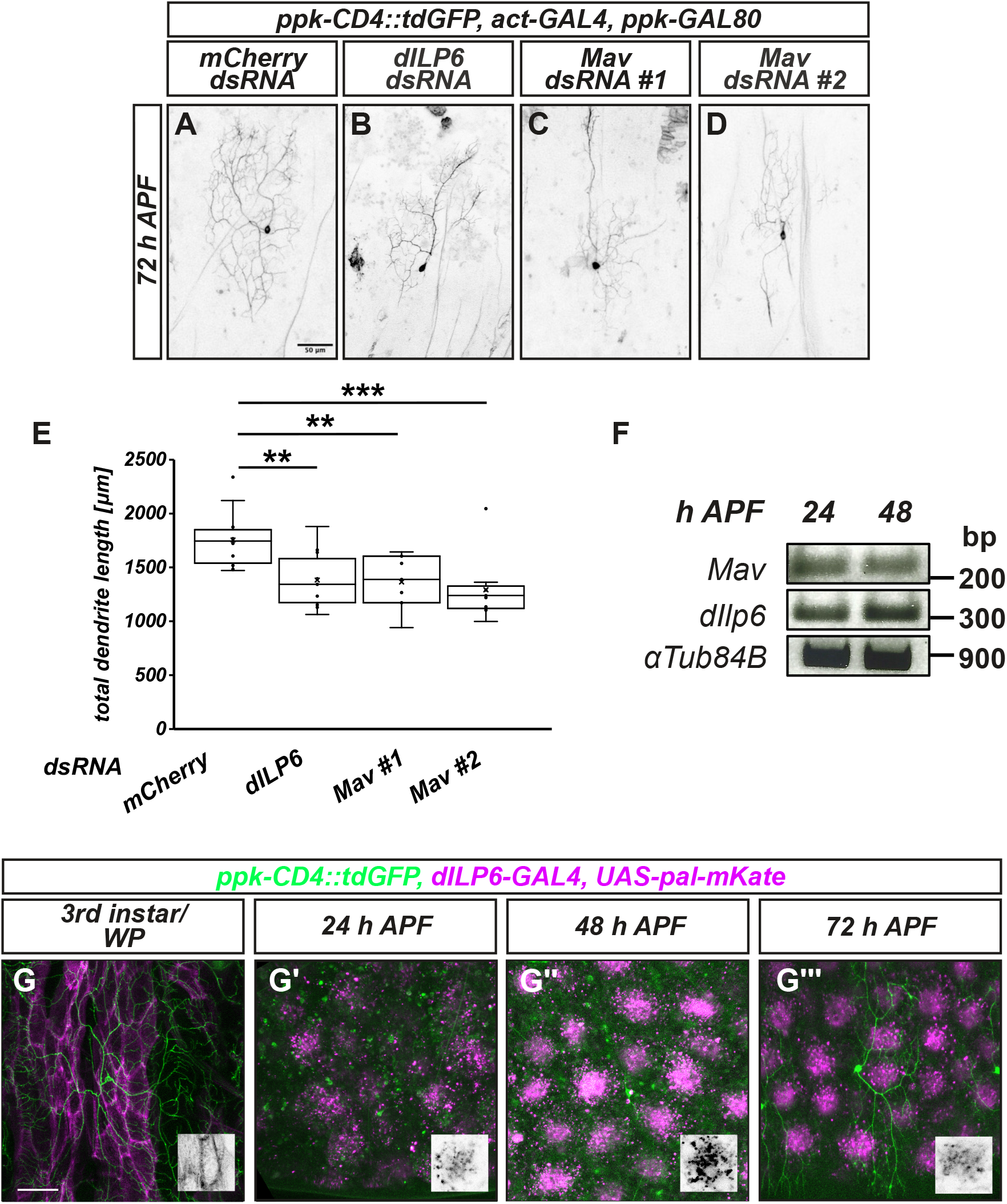
RTK ligands for dendrite regrowth are linked to the epidermis. **A** - **D** Non-cell autonomous knockdown of *dIlp6* or *Maverick* causes c4da neuron dendrite regrowth defects. dsRNA constructs were driven under *act5C-GAL4* in the presence of *ppk-GAL80* to suppress expression in c4da neurons. C4da neuron morphology was assessed at 72 h APF using *ppk-CD4::tdGFP*. **A** C4da neuron in a control animal expressing mCherry dsRNA. **B** C4da neuron in an animal expressing dIlp6 dsRNA. **C** C4da neuron in an animal expressing Mav dsRNA #1. **D** C4da neuron in an animal expressing Mav dsRNA #2. **E** Dendrite length in samples **A** - **D**. N=7 - 12, neurons, ** P<0.01, *** P<0.0001, Wilcoxon’s test. **F** RT-PCR analysis of dILP6 and Mav expression in pupal bodywalls at 24 or 48 h APF. **G** - **G’’’** Timecourse and pattern of *dILP6-GAL4* expression in the larval and pupal bodywall between the third instar larval stage and 72 h APF. Insets show single clusters of epidermal cells labeled by *dILP6-GAL4*. Scale bars in **A** and **G** are 50 μm.

Mav is known to be expressed in the larval epidermis (Hoyer et al., 2018). As we could not detect Mav in pupal samples using publicly available GFP-tagged transgenes, we assessed Mav mRNA expression in pupal abdominal bodywall preparations using RT-PCR and found that it was expressed at relatively constant levels at 24 and 48 h APF (Fig. 3 F). We next assessed dIlp6 expression and could also detect it via RT-PCR (Fig. 3 F). To assess dIlp6 expression in the abdominal body wall in more detail, we next used a GAL4 insertion in the *ILP6* gene to drive a membrane-bound red fluorescent protein, while labeling c4da neurons with *ppk-CD4::tdGFP*. *dILP6-GAL4* was clearly expressed in many LECs (Fig. 3 G). After metamorphic epidermal remodeling, *dILP6-GAL4* was expressed in regularly spaced foci of four to six histoblasts (Fig. 3 H-J, insets), with little or no expression in the histoblasts between them, potentially reminiscent of patterns caused by lateral inhibition. This expression pattern was already visible at 24 h APF (Fig. 3 H), when *dILP6-GAL4* activity was still relatively weak. *dILP6-GAL4* activity was strongest at 48 h APF (Fig. 3 I) and was still high at 72 h APF (Fig. 3 J).

Consistent with the idea that several insulin-like peptides might locally support c4da neuron dendrite regrowth, we found that *ILP2-GAL4*, but not *dILP3-GAL4* or *dILP5-GAL4*, was also expressed in epidermal cells. At the larval stage, it was active in very few LECs (Fig. S4 A). While *ILP2-GAL4* activity was low or absent at 24 and 48 h APF, where only cell debris was visible (Fig. S4 A’, A’’), it could be seen in many discrete histoblasts at 72 h APF (Fig. S4 A’’’). While we could not detect *dILP5-GAL4* activity in abdominal tissues (Fig. S4 C’ - C’’’), we noted that *dILP3-GAL4* was expressed in a structure reminiscent of muscle or gut at the larval stage (Fig. S4 B). Thus, our data suggest that dendrite regrowth is stimulated by Ret and InR ligands that are (sometimes dynamically) expressed in locally adjacent tissues in the pupal bodywall, especially in epidermal cells.

### LEC apoptosis is required for both c4da neuron dendrite pruning and regrowth

The above observations suggested that epidermal cells are functionally involved in dendrite regrowth. Such a link would be interesting since the epidermis is already important during c4da neuron dendrite pruning: morphogenetic movements in the epidermis help to break dendrites mechanically (Krämer et al., 2023), LECs have been proposed to send pruning-promoting signals to c4da neurons (Xu et al., 2026), and LECs act as phagocytes that remove dendrite debris after dendrite severing and fragmentation (Han et al., 2014). After dendrite regrowth, the adult c4da neuron dendrites are also in close contact with the epidermis (Yasunaga et al., 2010). LECs start to undergo apoptosis around 18 h APF, when histoblasts start to expand (Ninov et al., 2007). Given the close relationship between the two tissues, we wanted to know when epidermis remodeling occurs in relation to dendrite remodeling. To this end, we followed the progression of both c4da neuron dendrite remodeling (using *ppk-CD4::tdGFP*) and epidermis development (using *ECad::RFP*) together in the same animals. At the third instar larval stage, the epidermis was formed by large LECs, the histoblast nests were small, and c4da neurons had long and branched dendrites (Fig. 4 A, A’). At 24 h APF, condensed cells expressing *ECad::RFP* were frequently seen in the epidermis, and the area of the histoblast nests had significantly increased, indicating ongoing LEC apoptosis and histoblast expansion (Fig. 4 B). At this stage, all c4da neuron dendrites had been pruned (Fig. 4 B’). At 48 and 72 h APF, the epidermis completely consisted of histoblasts, and occasional ECad::RFP-positive blobs underneath the epidermal layer suggested that residual apoptotic LEC debris was being removed (Fig. 4 C, D). Consistent with our previous data (Sanal et al., 2023), c4da neuron dendrites had just started to regrow at 48 h APF (Fig. 4C’), and had regrown long and branched dendrites at 72 h APF (Fig. 4 D’). Thus, both LEC apoptosis and histoblast expansion largely occur during a time window when c4da neuron dendrite pruning is finished and dendrite regrowth has not yet started.

**Figure 4.**
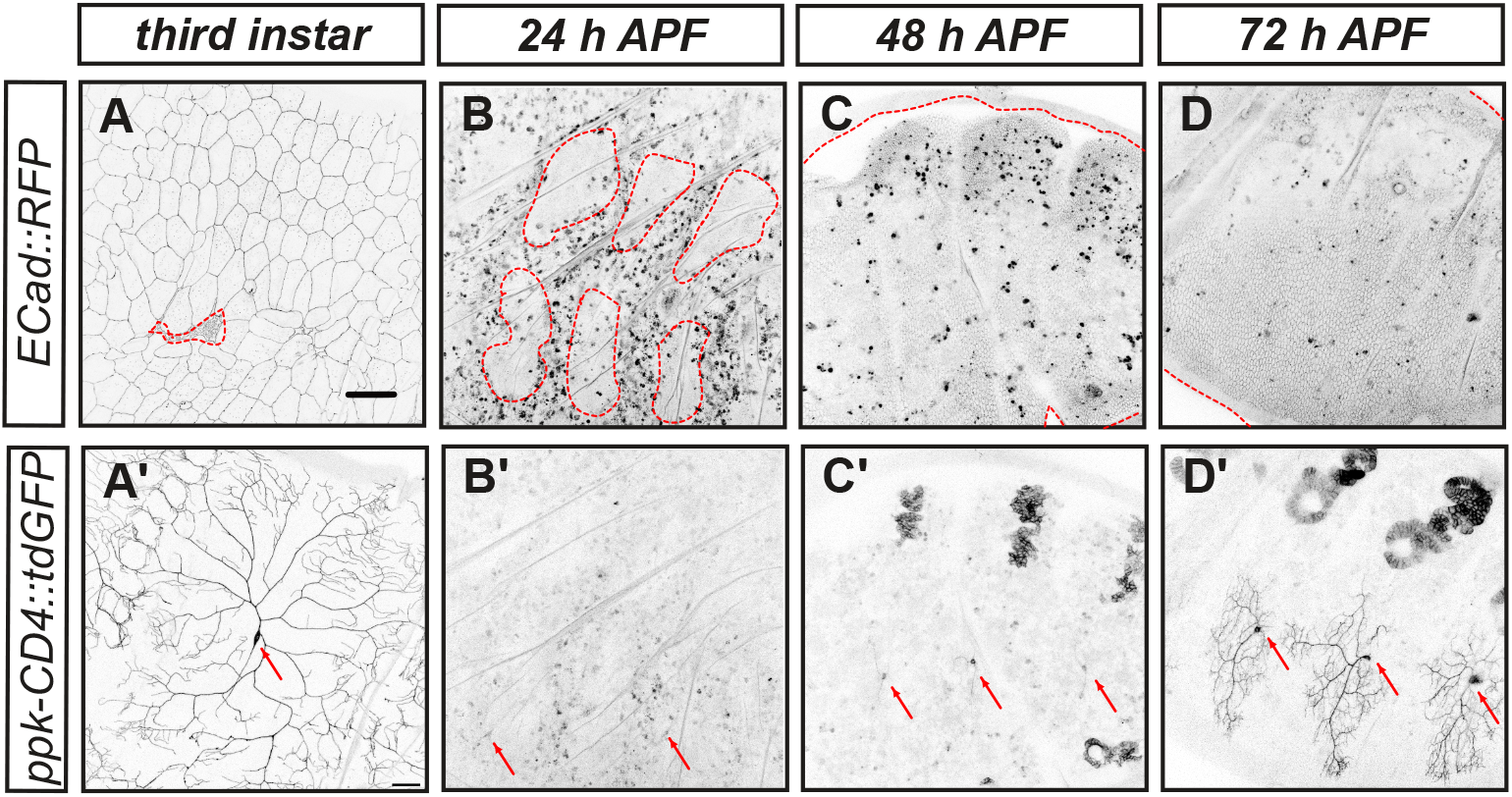
Epidermis remodeling occurs after c4da neuron dendrite pruning and before dendrite regrowth. **A** - **D’** Epidermal cells (**A** - **D**) and c4da neurons (**A’** - **D’**) were visualized at the indicated timepoints using ECad::3xRFP and ppk-Cd4::tdGFP, respectively. **A**, **A’** Third instar larval stage. **B**, **B’** 24 h APF. **C**, **C’** 48 h APF. **D**, **D’** 72 h APF. Histoblast nests and areas covered by histoblasts are demarkated by dashed red lines in **A** - **D**, positions of c4da neuron cell bodies in **A’** - **D’** are indicated by arrows. The scale bar in **A** is 50 μm.

To test for a direct link between the two processes, we next sought ways to prevent epidermis remodeling. Many manipulations preventing LEC remodeling result in early pupal death with strong defects in pupal ecdysis, thus preventing analysis past 24 h APF, but we found that expression of the caspase inhibitor p35 in LECs under the *A58-GAL4* driver resulted in animals that developed at least until 72 h APF, even though defects in head eversion were frequently seen. In these animals, LEC morphology and histoblast nests looked normal at the third instar larval stage (Fig. 5 A). At 24 h APF, condensed ECad-positive structures could be seen between the LECs, suggesting that some LECs still initiated or underwent apoptosis, but the histoblast nests in these animals had not expanded significantly from their larval size and were smaller than in control animals (Fig. 5 B, E). At 48 and 72 h APF, the histoblasts nests had still not expanded significantly (apart from occasional escapers), and the epidermis still consisted of surviving LECs (Fig. 5 C-E), indicating a major delay in epidermis remodeling. While c4da neuron morphology was normal at the third instar (Fig. 5 F), we observed strong c4da neuron dendrite pruning defects at 24 h APF, where over 60% of neurons still had dendrites attached to the soma in these animals (Fig. 5 G, M). At the 48 h APF timepoint, more than half of the c4da neurons still had larval dendrites (Fig 5 H, M). Only at 72 h APF had most c4da neurons pruned their larval dendrites, even though larval dendrite debris could still be detected (Fig. 5 I, M).

**Figure 5.**
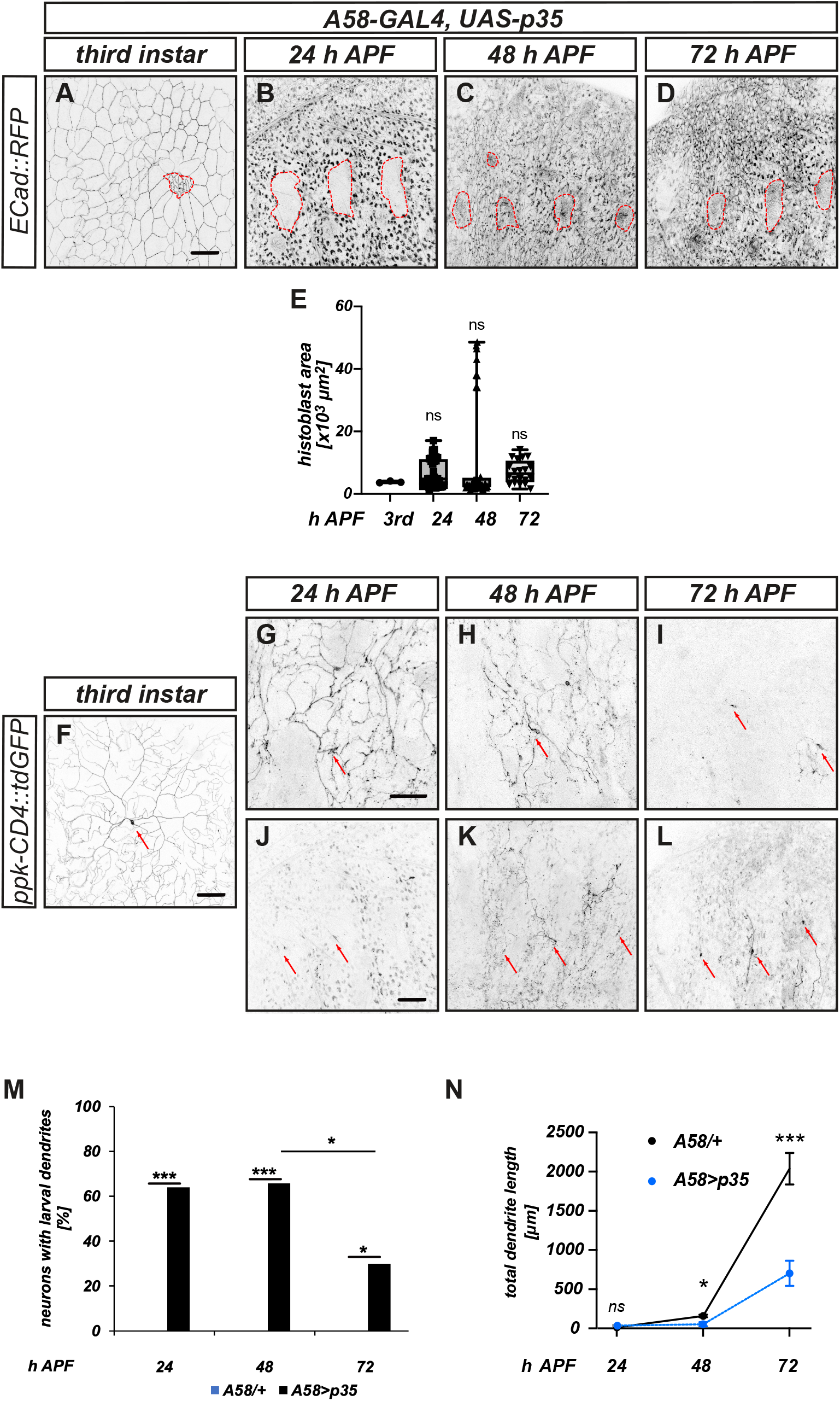
Epidermis remodeling is required for c4da neuron dendrite pruning and regrowth. **A** - **D** Epidermal cells were visualized with ECad::RFP in animals expressing the caspase inhibitor p35 under the LEC driver A58-GAL4 at the indicated times after puparium formation. Histoblast nests are marked by dashed red lines. **A** Third instar larval stage. **B** 24 h APF. **C** 48 h APF. **D** 72 h APF. **E** Quantification of epidermis area covered by histoblasts in **A** - **D**. N= 3, 36, 37, 20, values are mean +/- s. d., n. s., not significant, Mann Whitney test. **F** C4da neuron labeled by *ppk-CD4::tdGFP* in a third instar larva expressing p35 in LECs. Panels **G** - **I** show images of pupal c4da neurons in these animals that did not or only partially undergo dendrite pruning. **G** C4da neuron in 24 h APF animal. **H** C4da neuron in 48 h APF animal. **I** C4da neuron in 72 h APF animal. Panels **J** - **L** show images of pupal c4da neurons in these animals that did undergo dendrite pruning. **J** C4da neuron in 24 h APF animal. **K** C4da neuron in 48 h APF animal. **L** C4da neuron in 72 h APF animal. **M** Percentage of neurons with unpruned (attached) larval dendrites at 24, 48 and 72 h APF. Blue bars: A58/+ control animals (N=13, 22, 15), black bars: A58>p35 animals (N=52, 34, 20), n. s., not significant, * P<0.05, *** P<0.0005, Fisher’s exact test. **N** Quantification of the length of regrown dendrites, comparing the subset of c4da neurons in A58>p35 animals (in **J** - **L**) that did undergo pruning with c4da neurons in control animals only carrying A58 see **Fig. 4 B’ - D’**). N=10-12 neurons each, values are mean +/- s. d., n. s., not significant, * P<0.05, *** P<0.001, Mann Whitney test. Scale bars in **A**, **F**, **G**, **J** are 50 μm.

In order to assess whether inhibition of epidermis remodeling also affects dendrite regrowth, we next measured the length of newly grown dendrites in those c4da neurons in which dendrite pruning had occured. As expected, no new dendrites could be observed in these animals at 24 h APF (Fig. 5 J), similar to what was seen in control animals (Fig. 5 J, N). At the 48 h APF timepoint, when c4da neurons in control animals had just started dendrite regrowth (e. g., Fig. 4 C’), no regrown dendrites could be observed in these animals either (Fig. 5 K, N), and at 72 h APF, the difference in dendrite length was highly significant, as control neurons had long and branched dendrites (e. g., Fig. 4 D’), whereas those in the animals with inhibited epidermis remodeling were still absent or very short (Fig. 5 L, N). Thus, LEC apoptosis and epidermis remodeling are not only important for c4da neuron dendrite pruning, but they are also also required for dendrite regrowth.

### Dendrite remodeling may be evolutionarily linked to epidermis remodeling

As both larval and adult c4da neuron dendrites are in tight contact with epidermal cells, which act as their growth and adhesion substrates (Yasunaga et al., 2010; Han et al., 2012; Kim et al., 2012), a plausible explanation for the observed link between dendrite remodeling and epidermis remodeling is that c4da neuron dendrite pruning and regrowth are necessary because the larval dendrites cannot be maintained without their adhesive substrate during epidermis remodeling. In such a model, epidermal (re-)expression of RTK ligands, and potentially the cessation of expansion-associated tissue movements, could signal the reemergence of a stable growth substrate to induce dendrite regrowth. This model is testable because not all holometabolic insects undergo epidermis remodeling. In Drosophila, LECs are highly polyploid so LEC growth can keep up with larval body growth, and their metamorphic apoptosis is likely necessary because these cells cannot undergo cell division and other morphogenetic changes to adapt to adult skin morphology. In contrast, the epidermis of other insects, including other dipteres such as *Aedes aegypti* mosquitos, consists of diploid or only tetraploid epidermal cells throughout development, such that these cells can still divide during metamorphosis and do not undergo apoptosis (Trager, 1937; Risler, 1959). We were therefore curious to know whether *Aedes* peripheral sensory neurons undergo dendrite pruning. Antibodies against horseradish peroxidase (HRP) stain peripheral neurons in a wide range of insect species (Jan & Jan, 1982). We used this to visualize peripheral bodywall neurons in filets of mosquito fourth instar larvae and 24 h APF pupae. This showed a similar amount of neurites for both larval and pupal PNS neurons (Fig. 6 A, A’, B, B’). For comparison, we also visualized all Drosophila PNS neurons at the third instar or 24 h APF by expressing tdtomato under the control of a panneuronal *nsyb-GAL4* driver. As expected, much fewer neurites could be seen in the pupal bodywall compared to the larval one (Fig. 6 C, D). Thus, PNS neuron dendrite remodeling may also be linked epidermis remodeling during evolution.

**Figure 6.**
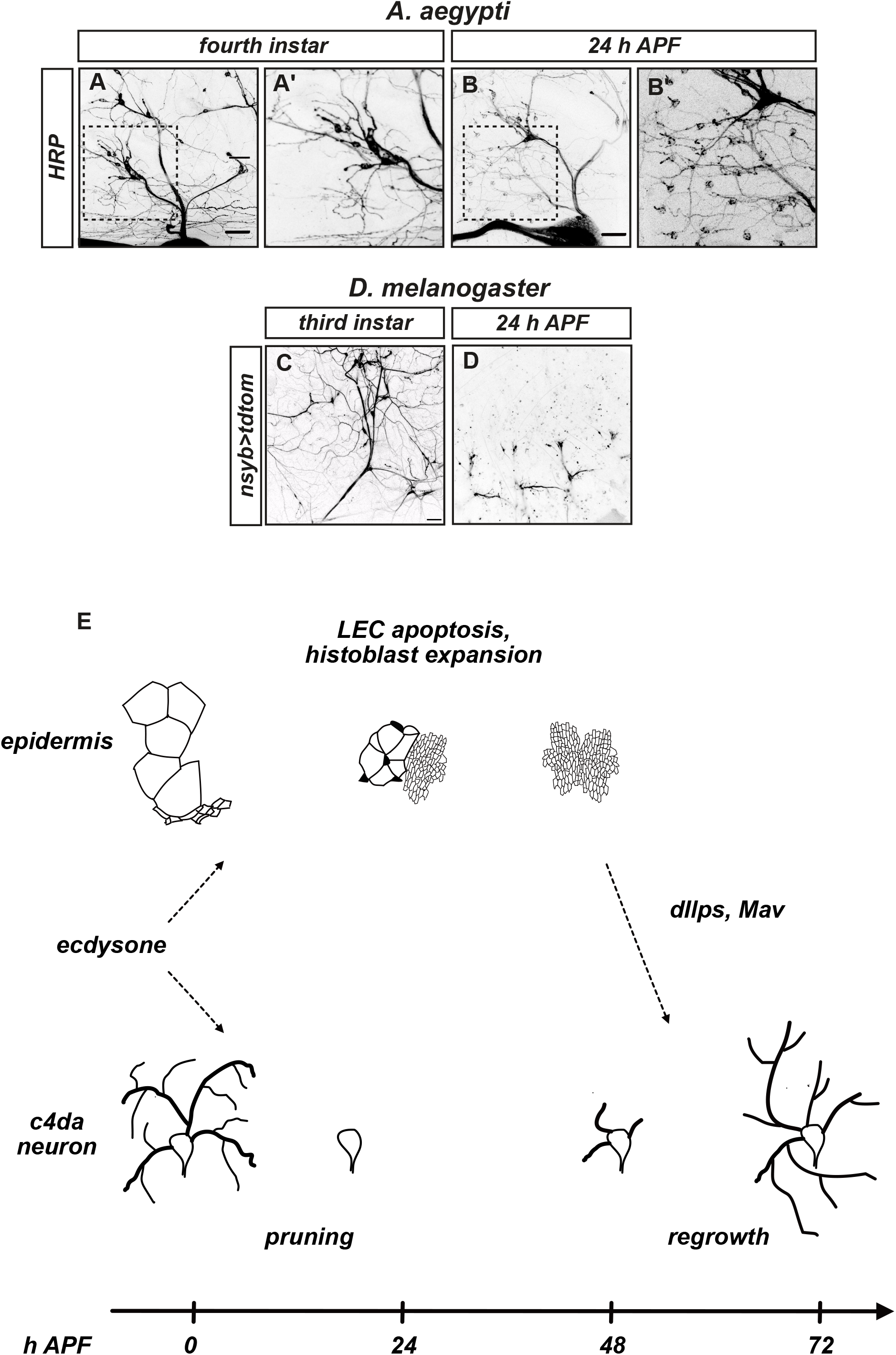
No evidence for dendrite remodeling during metamorphosis in *Aedes aegypti*. **A** - **B’** *Aedes aegypti* peripheral neurons and neurites in the ventral bodywall were visualized by immunofluorescence using HRP antibodies at the fourth instar stage (**A**) or at 24 h APF (**B**). Panels **A’**, **B’** show enlarged images of the boxed areas in **A**, **B**. **C**, **D** Peripheral neurons and neurites in the Drosophila bodywall were visualized by tdtomato expression under the pan-neuronal driver R57C10(nsyb)-GAL4 at the third instar stage (**C**) or at 24 h APF (**D**). **E** Hypothetical model for the coordination between c4da neuron dendrite pruning and regrowth via the epidermis. Scale bars are 10 μm in **A** and **D**.

## Discussion

In this work, we assessed signals leading to the initiation of c4da neuron dendrite regrowth after pruning. Our data indicate that dendrite regrowth depends on the two RTKs InR and Ret, whose ligands are derived locally, and at least in part from the epidermal cells that form the adult epidermis. As larval epidermal cells are replaced by histoblasts during metamorphosis, we tested whether epidermis remodeling is also required for dendrite remodeling. Our results show that inhibition of epidermal cell remodeling causes strong defects in both dendrite pruning and dendrite regrowth, indicative of tight coordination etween the two processes. A plausible model would therefore be that epidermal (re-)expression of RTK ligands, and potentially the cessation of expansion-associated tissue movements, could signal the reemergence of a stable growth substrate to induce dendrite regrowth.

Epidermal cells would thus be involved in the coordination between dendrite pruning and regrowth. Epidermis remodeling - both LEC apoptosis and histoblast expansion - is induced by ecdysone at the onset of metamorphosis (Ninov et al., 2007), similarly to dendrite pruning. While c4da neuron dendrite regrowth does not depend on cell autonomous ecdysone signaling (Fig. 1), it is likely a late consequence of this ecdysone signaling via a different cell type (Fig. 6 E). In our model, ecdysone first simultaneously induces dendrite pruning, LEC apoptosis and histoblast expansion. Once the histoblasts have expanded and form the new epidermis, they secrete RTK ligands to induce dendrite regrowth. Ecdysone would thus regulate both dendrite pruning and regrowth via an incoherent feed-forward loop (Mangan and Alon, 2003), where a direct and fast, cell autonomous response to ecdysone in the c4da neurons induces pruning, but a slow and indirect, non-cell autonomous response to ecdysone via the epidermis induces the opposite response, dendrite regrowth (Fig. 6 E).

In further support of a link between epidermis remodeling and dendrite remodeling, Aedes mosquitos lacking epidermis remodeling also apparently do not appear to have sensory neuron dendrite remodeling. Interestingly, metamorphic remodeling appears to be much more prominent in Drosophila tissues, where the duration of both the larval stage and metamorphosis are approximately five days, compared to Aedes, where the larval stage is eight days long, but metamorphosis only takes two days. Thus, metamorphic apoptosis of polyploid LECs acompanied by dendrite remodeling may be an evolutionary mechanism to enable a shorter larval stage, at the expense of a longer pupal one.

Lastly, it was surprising that apoptosis inhibition in LECs caused strong dendrite pruning defects (Fig. 5). We think that preventing LEC apoptosis and histoblast expansion could interfere with the morphogenetic movements that break the c4da neuron dendrites in their proximal regions (Krämer et al., 2023). Taken together, our data reveal a tight coordination between sensory neuron and epidermis development during metamorphosis.

## Materials and Methods

### Fly Strains

For expression in c4da neurons, we used *ppk-GAL4* insertions on the second and third chromosomes (Grueber et al., 2007) driving UAS-CD8::GFP or UAS-tdtomato (Han et al., 2014) or *ppk-shine-GAL4* (this study) for inducible expression. Larval epidermal cells were manipulated using A58-GAL4 (Galko and Krasnow, 2004). For ubiquitous expression with exception of c4da neurons, we used *act5C-GAL4* (BL4414) recombined with *ppk-GAL80* (Yang et al., 2009), and *GAL4(R57C10)* (BL39171) for panneuronal expression. Expression patterns of insulin-like peptides were assessed with *dILP2-GAL4* (BL37516), *dILP3-GAL4* (BL52660), *dILP5-GAL4* (BL66007) and *dILP6-GAL4* (Kyoto 103877). Epidermal cells were labeled with ECad::RFP (Pineiro et al., 2017), and c4da neurons independently of GAL4 with *ppk-CD4::tdGFP* (Han et al., 2011). The Ret mutant used was *RetΔ/Df BSC312* (Hoyer et al., 2018), and Ret expression was assessed using Ret^MI07200^ (BL43099). UAS lines were UAS-EcR^DN^ (BL 6872), UAS-pal-mKate (gift from S. Luschnig), UAS-Ret (Hoyer et al., 2018), UAS-PI3K CAAX (BL8294), UAS-p35 (BL5073). All dsRNAs were coexpressed with UAS-dcr2 (Dietzl et al., 2007).

### List of dsRNA lines

|  |  |
| --- | --- |
| Orco (control dsRNA) | BL31278 |
| mCherry (control) | BL35785 |
| Ret #1 | BL25948 |
| Ret #2 | BL55872 |
| InR | VDRC991 |
| Alk | BL27518 |
| Alk | VDRC11446 |
| Eph | BL28511 |
| Lgr3 | BL55910 |
| NF1 | BL29845 |
| Nrk | BL55184 |
| Nrk | BL56936 |
| Nrk | VDRC36282 |
| Otk | BL25970 |
| Ror | NIG-Fly 4926R-1 |
| Ror | VDRC29930 |
| Toll-6 | BL56048 |
| Toll-7 | BL30488 |
| activin $\beta$ | BL42493 |
| daw | BL34974 |
| daw | BL50911 |
| gbb | BL34898 |
| Ilp1 | BL32861 |
| Ilp2 | BL31068 |
| Ilp3 | BL33681 |
| Ilp5 | BL33683 |
| Ilp6 | BL31379 |
| Ilp7 | BL32862 |
| Ilp8 | BL80436 |
| Ilp8 | BL82979 |
| mav | BL34650 |
| mav | BL36809 |
| myoglianin | BL31114 |
| wingless | BL31310 |
| wingless | BL33902 |
| wnt2 | BL29441 |
| wnt2 | BL44271 |
| wnt5 | BL28534 |
| wntless | VDRC5214 |

### Cloning and transgenes

DBD-nMag^high1^ and pMag^high1^-AD inserts were generated by PCR from corresponding plasmids (di Pietro et al., 2021) and cloned into pDEST APHIH (containing the *ppk* promotor, gift from Chun Han) using the TOPO/TA system. The resulting *ppk-DBD-nMag^high1^* and *ppk-pMag^high1^-AD1* plasmids were inserted into the VK37 (second chromosome) and 86Fb (third chromosome) insertion sites, respectively, and crossed together to yield a *ppk-shine-GAL4* line.

### Dissection, Microscopy and Live imaging

For analyses of dendrite and epidermis morphology at different timepoints during larval or pupal stages, larvae or pupae dissected out of the pupal case were mounted in PBS/90% glycerol. Lateroventral v’ada c4da neurons in segments A3 - A5 were imaged live on a Zeiss LSM710 confocal microscope with a 20x Plan Apochromat water objective (1.0 NA). All processing was done in Fiji (Schindelin *et al*, 2012).

### RT-PCR

Ten to twelve pupae each at the 24, 48, and 72 h APF timepoints were dissected out of the pupal case, and total RNA was extracted from abdomens without gut using Trizol (Invitrogen cat. # 15596026). cDNA was generated using the Superscript III kit (Invitrogen cat. # 18080044) according to the manufacturer’s instructions. PCR primers were CGGCCTAGAAACTAAGAGGC and GTTGTTAAAATAGCTGTTGCC for Maverick, ATGGTTCTCAAAGTGCCGAC and CTATCCTTTGCAGTACTGCAAG for dILP6, and ACAACGAGGCTATCTACGACA and TTTTCAGTGTTGCAGTGAATTT for αTub84B.

### Immunofluorescence

Mosquito eggs (strain “Monheim”, Envu Germany, courtesy of J. Egger, Envu) were hatched in ddH_2_O supplemented with fish food at 25°C. Appropriately staged late larvae and early pupae were collected for dissection, and residual cultures were inactivated by adding acetic acid to 5% v/v. Larvae and pupae were dissected dorsally into body wall filets in ice-cold PBS and fixed in PBS containing 4% formaldehyde and 2% Triton X-100 over night at 4°C, washed three times in PBS with 1 % Triton X-100 and blocked in PBS with 1 % Triton X-100 and 10% goat serum for one hour. Filets were incubated with Cy3-conjugated antibodies against HRP (1:1000) in blocking solution for 48 hours and washed in PBS with 1 % Triton X-100 three times before mounting.

### Quantification and statistical analyses

Phenotypic penetrance was assessed by counting the number of neurons with dendrites still attached to the soma. Here, significance was determined using a categorical two-tailed Fisher’s exact test. Total dendrite length was determined using the Fiji plugin NeuronJ and compared using the Mann Whitney test. Prism software was used for all statistical analyses.

## Supporting information

Sanal et al., 2026 Supplementary Figures

## Acknowledgments

We thank F. di Pietro and Y. Bellaiche for shine-GAL4 plasmids and flies, J. Egger, S. Horstmann and F. Käding (Envu Germany, Monheim) for Aedes egg collections, P. Soba, C. Klämbt, C. Han and S. Luschnig for fly lines, Lorena Keding for early contributions to the project, and the Bloomington and VDRC stock centers for fly lines. NS was a member of the CiM-IMPRS graduate school. This work was supported by DFG grants RU1673/4-1 and RU1673/6-1 to SR, SFB1348 grant B04 to SR, and support by the Forschungsförderungsfond of the Faculty of Biology at the University of Münster to SR. The authors declare no competing financial interests.

## Author Contributions

NS, UG, and SR performed experiments, UG and SR generated reagents. NS and SR contributed to experimental design. NS and SR analyzed data, and SR wrote the manuscript.

## Supplementary Figure legends

**Figure S1. RNAi of RTKs and effect on dendrite regrowth.** The indicated dsRNA lines were crossed to *2xppk-GAL4, UAS-CD8::GFP, UAS-dcr2,* and dendrite morphology was assessed at 72 h APF. **A** Orco control dsRNA. **B** Alk dsRNA #1. **C** Alk dsRNA #2. **D** Eph dsRNA. **E** Lgr3 dsRNA. **F** NF1 dsRNA. **G** Nrk dsRNA #1. **H** Nrk dsRNA #2. **I** Nrk dsRNA #3. **J** Otk dsRNA. **K** Ror dsRNA #1. **L** Ror dsRNA #2. **M** Toll6 dsRNA. **N** Toll7 dsRNA. The scale bar in A is 50 μm.

**Figure S2. RET expression in pupal c4da neurons. A** - **A’’’** *Ret^MI07200^*expression at the indicated timepoints between the white pupal stage and 72 h APF. The scale bar in **A** is 20 μm.

**Figure S3. RNAi of candidate RTK ligands and effect on dendrite regrowth.** The indicated dsRNA lines were crossed to *act-GAL4, ppk-GAL80, ppk-CD4::tdGFP* and dendrite morphology was assessed at 72 h APF. **A** mCherry control dsRNA. **B** Ilp1 dsRNA. **C** Ilp2 dsRNA. **D** Ilp3 dsRNA. **E** Ilp5 dsRNA. **F** Ilp7 dsRNA. **G** Ilp8 dsRNA #1. **H** Ilp8 dsRNA #2. **I** Wnt2 dsRNA. **J** Wnt5 dsRNA. The scale bar in A is 50 μm.

**Figure S4. *dILP-GAL4* expression in the larval and pupal epidermis. A** - **A’’’** Timecourse and pattern of *ILP2-GAL4* expression in the larval and pupal bodywall between the third instar larval stage and 72 h APF. Insets show single clusters of epidermal cells labeled by *dILP2-GAL4*. **B** - **B’’** Timecourse of *ILP3-GAL4* expression in larval and pupal peripheral tissues between the third instar larval stage and 72 h APF. **C** - **C’’’** Timecourse of *ILP5-GAL4* expression in the larval and pupal bodywall between the white pupal stage and 72 h APF. All scale bars are 50 μm.

## References

Arensburger, P., Megy, K., Waterhouse, R.M., Abrudan, J., Amedeo, P., Antelo, B., Bartholomay, L., Bidwell, S., Caler, E., Camara, F., et al., 2010. Sequencing of Culex quinquefasciatus establishes a platform for mosquito comparative genomics. Science 330, 86–88.

Davies, M., Sanal, N., Wolterhoff, N., Gigengack, U., Shen, Y., Hahn, I., Rumpf, S. 2025. Spectraplakin cooperates with noncentrosomal microtubule regulators to orient dendritic microtubules in Drosophila. eLife doi:10.7554/eLife.108055.1

Dietzl, G., Chen, D., Schnorrer, F., Su, K.-C., Barinova, Y., Fellner, M., Gasser, B., Kinsey, K., Oppel, S., Scheiblauer, S., et al., 2007. A genome-wide transgenic RNAi library for conditional gene inactivation in Drosophila. Nature 448, 151–156.

di Pietro, F., Herszterg, S., Huang, A., Bosveld, F., Alexandre, C., Sancéré, L., Pelletier, S., Joudat, A., Kapoor, V., Vincent, J.P., et al., 2021. Rapid and robust optogenetic control of gene expression in Drosophila. Dev. Cell 56, 3393–3404.e7.

Frommeyer, S.M., Gigengack, U., Rode, S., Davies, M., Wolterhoff, N., Rumpf, S., 2026. The mRNA export factor UAP56 is required for dendrite and synapse pruning via actin regulation in Drosophila. J Cell Sci. 139, jcs264770.

Galko, M.J., Krasnow, M.A., 2004. Cellular and genetic analysis of wound healing in Drosophila larvae. PLoS Biol 2, E239.

Grueber, W.B., Ye, B., Yang, C.-H., Younger, S., Borden, K., Jan, L.Y., Jan, Y.-N., 2007. Projections of Drosophila multidendritic neurons in the central nervous system: links with peripheral dendrite morphology. Development 134, 55–64.

Han, C., Jan, L.Y., Jan, Y.N., 2011. Enhancer-driven membrane markers for analysis of nonautonomous mechanisms reveal neuron-glia interactions in Drosophila. Proc. Natl. Acad. Sci. U.S.A. 108, 9673–8.

Han, C., Wang, D., Soba, P., Zhu, S., Lin, X., Jan, L.Y., Jan, Y.-N., 2012. Integrins regulate repulsion-mediated dendritic patterning of drosophila sensory neurons by restricting dendrites in a 2D space. Neuron 73, 64–78.

Han, C., Song, Y., Xiao, H., Wang, D., Franc, N.C., Jan, L.Y., Jan, Y.-N., 2014. Epidermal cells are the primary phagocytes in the fragmentation and clearance of degenerating dendrites in Drosophila. Neuron 81, 544–560.

Hoyer, N., Zielke, P., Hu, C., Petersen, M., Sauter, K., Scharrenberg, R., Peng, Y., Kim, C.C., Han, C., Parrish, J.Z., et al., 2018. Ret and Substrate-Derived TGF-β Maverick Regulate Space-Filling Dendrite Growth in Drosophila Sensory Neurons. Cell Rep. 24, 2261–2272.e5.

Jan, L.Y., Jan, Y.N., 1982. Antibodies to horseradish peroxidase as specific neuronal markers in Drosophila and in grasshopper embryos. Proc. Natl. Acad. Sci. U.S.A. 79, 2700–4.

Johnson, M.B., Stevens, B., 2018. Pruning hypothesis comes of age. Nature, 554, 438–439.

Kim, D, Dressler, G.R., 2007. PTEN modulates GDNF/RET mediated chemotaxis and branching morphogenesis in the developing kidney. Dev. Biol. 307, 90–99.

Kim, M.E., Shrestha, B.R., Blazeski, R., Mason, C.A., Grueber, W.B., 2012. Integrins establish dendrite-substrate relationships that promote dendritic self-avoidance and patterning in drosophila sensory neurons. Neuron 73, 79–91.

Kirilly, D., Gu, Y., Huang, Y., Wu, Z., Bashirullah, A., Low, B.C., Kolodkin, A.L., Wang, H., Yu, F., 2009. A genetic pathway composed of Sox14 and Mical governs severing of dendrites during pruning. Nat. Neurosci. 12, 1497–1505.

Krämer, R., Wolterhoff, N., Galic, M., Rumpf, S., 2023. Developmental pruning of sensory neurites by mechanical tearing in Drosophila. J. Cell Biol. 222.

Kuo, C.T., Jan, L.Y., Jan, Y.N., 2005. Dendrite-specific remodeling of Drosophila sensory neurons requires matrix metalloproteases, ubiquitin-proteasome, and ecdysone signaling. Proc. Natl. Acad. Sci. U.S.A. 102, 15230–15235.

Lee T, Marticke S, Sung C, Robinow S, Luo L., 2000. Cell-autonomous requirement of the USP/EcR-B ecdysone receptor for mushroom body neuronal remodeling in Drosophila. Neuron 28, 807–18.

Luo, W., Enomoto, H., Rice, F.L., Milbrandt, J., Ginty, D.D., 2009. Molecular identification of rapidly adapting mechanoreceptors and their developmental dependence on ret signaling. Neuron 64, 841–56.

Mangan, S., Alon, U., 2003. Structure and function of the feed-forward loop network motif. Proc Natl Acad Sci U S A. 100, 11980–5.

Michel, M., Dahmann C., 2020. Tissue mechanical properties modulate cell extrusion in the Drosophila abdominal epidermis. Development 147, dev179606.

Marzano, M., Herzmann, S., Elsbroek, L., Sanal, N., Tarbashevich, K., Raz, E., Krahn, M.P., Rumpf, S., 2021. AMPK adapts metabolism to developmental energy requirement during dendrite pruning in Drosophila. Cell Rep. 37, 110024.

Myers, L., Perera, H., Alvarado, M.G., Kidd, T., 2018. The *Drosophila Ret* gene functions in the stomatogastric nervous system with the Maverick TGFβ ligand and the *Gfrl* co-receptor. Development 145, dev157446.

Ninov, N., Chiarelli, D.A., Martin-Blanco, E., 2007. Extrinsic and intrinsic mechanisms directing epithelial cell sheet replacement during Drosophila metamorphosis. Development 134, 367–79.

Perea, D., Guiu, J., Hudry, B., Konstantinidou, C., Milona, A., Hadjieconomou, D., Carroll, T., Hoyer, N., Natarajan, D., Kallijärvi, J., et al., 2017. Ret receptor tyrosine kinase sustains proliferation and tissue maturation in intestinal epithelia. EMBO J. 36, 3029–3045.

Pinheiro, D., Hannezo, E., Herszterg, S., Bosveld, F., Gaugue, I., Balakireva, M., Wang, Z., Cristo, I., Rigaud, S. U., Markova, O. et al., 2017. Transmission of cytokinesis forces via E-cadherin dilution and actomyosin flows. Nature 545, 103–107

Rabinovich D, Yaniv SP, Alyagor I, Schuldiner O., 2016. Nitric Oxide as a Switching Mechanism between Axon Degeneration and Regrowth during Developmental Remodeling. Cell 164, 170–182.

Riccomagno, M.M., Kolodkin, A.L., 2015. Sculpting neural circuits by axon and dendrite pruning. Annu. Rev. Cell Dev. Biol. 31, 779–805.

Risler, H., 1959. POLYPLOIDIE UND SOMATISCHE REDUKTION IN DER LARVENEPIDERMIS VON AEDES AEGYPTI L. (CULICIDAE). Chromosoma 10, 184–209

Rui, M., 2024. Recent progress in dendritic pruning of *Drosophila* C4da sensory neurons. Open Biol. 14, 240059.

Rumpf, S., Rode, S., Herzmann, S., Krumkamp, R. 2017. Mechanisms of Neurite Pruning. Neuroforum 23, 19–25.

Rumpf, S., Wolterhoff, N., Herzmann, S., 2019. Functions of Microtubule Disassembly during Neurite Pruning. Trends Cell Biol. 29, 291–297.

Sanal, N., Keding, L., Gigengack, U., Michalke, E., Rumpf, S., 2023. TORC1 regulation of dendrite regrowth after pruning is linked to actin and exocytosis. PLoS Genet. 19, e1010526.

Schindelin, J., Arganda-Carreras, I., Frise, E., Kaynig, V., Longair, M., Pietzsch, T., Preibisch, S., Rueden, C., Saalfeld, S., Schmid, B., et al., 2012. Fiji: an open-source platform for biological-image analysis. Nat. Methods 9, 676–682.

Scott-Hewitt, N., Perrucci, F., Morini, R., Erreni, M., Mahoney, M., Witkowska, A., Carey, A., Faggiani, E., Schuetz, L.T., Mason, S., et al., 2020. Local externalization of phosphatidylserine mediates developmental synaptic pruning by microglia. EMBO J. 39, e105380.

Trager, W., 1937. CELL SIZE IN RELATION TO THE GROWTH AND METAMORPHOSIS OF THE MOSQUITO, AEDES AEGYPTI. J. Exp. Zool. 3, 467– 489.

Williams, D.W., Truman, J.W., 2005. Cellular mechanisms of dendrite pruning in Drosophila: insights from in vivo time-lapse of remodeling dendritic arborizing sensory neurons. Development 132, 3631–3642.

Wong, J.J., Li, S., Lim, E.K., Wang, Y., Wang, C., Zhang, H., Kirilly, D., Wu, C., Liou, Y.C., Wang, H., et al., 2013. A Cullin1-based SCF E3 ubiquitin ligase targets the InR/PI3K/TOR pathway to regulate neuronal pruning. PLoS Biol. 11, e1001657.

Xu W, Wang S, Lv X, Long K, Mo K, Rui M., 2026. Patronin facilitates neurite remodeling via epithelial-to-neuronal signaling. Cell Commun. Signal. doi: 10.1186/s12964-026-03026-4.

Yang, C.-H., Rumpf, S., Xiang, Y., Gordon, M.D., Song, W., Jan, L.Y., Jan, Y.-N., 2009. Control of the postmating behavioral switch in Drosophila females by internal sensory neurons. Neuron 61, 519–526.

Yaniv, S.P., Issman-Zecharya, N., Oren-Suissa, M., Podbilewicz, B., Schuldiner, O., 2012. Axon regrowth during development and regeneration following injury share molecular mechanisms. Curr. Biol. 22,1774–82.

Yasunaga, K., Kanamori, T., Morikawa, R., Suzuki, E., Emoto, K., 2010. Dendrite reshaping of adult Drosophila sensory neurons requires matrix metalloproteinase-mediated modification of the basement membranes. Dev. Cell 18, 621–32.

Yin, C., Morita, T., Parrish, J.Z., 2024. A cell atlas of the larval *Aedes aegypti* ventral nerve cord. Neural Dev. 19, 2.

Yoong, L.F., Lim, H.K., Tran, H., Lackner, S., Zheng, Z., Hong, P., Moore, A.W., 2020. Atypical Myosin tunes dendrite arbor subdivision. Neuron 106, 452–467.e8.

Yuh Chew, L., He, J., Wong, J.J.L., Li, S., Yu, F., 2022. AMPK activates the Nrf2-Keap1 pathway to govern dendrite pruning via the insulin pathway in Drosophila. Development 149, dev200536.

