## Supplementary figures and images for "RTK signaling and epidermis remodeling instruct dendrite regrowth after pruning in the fly PNS"

### Sanal et al., 2026 Supplementary Figures

#### Sanal Figure S1

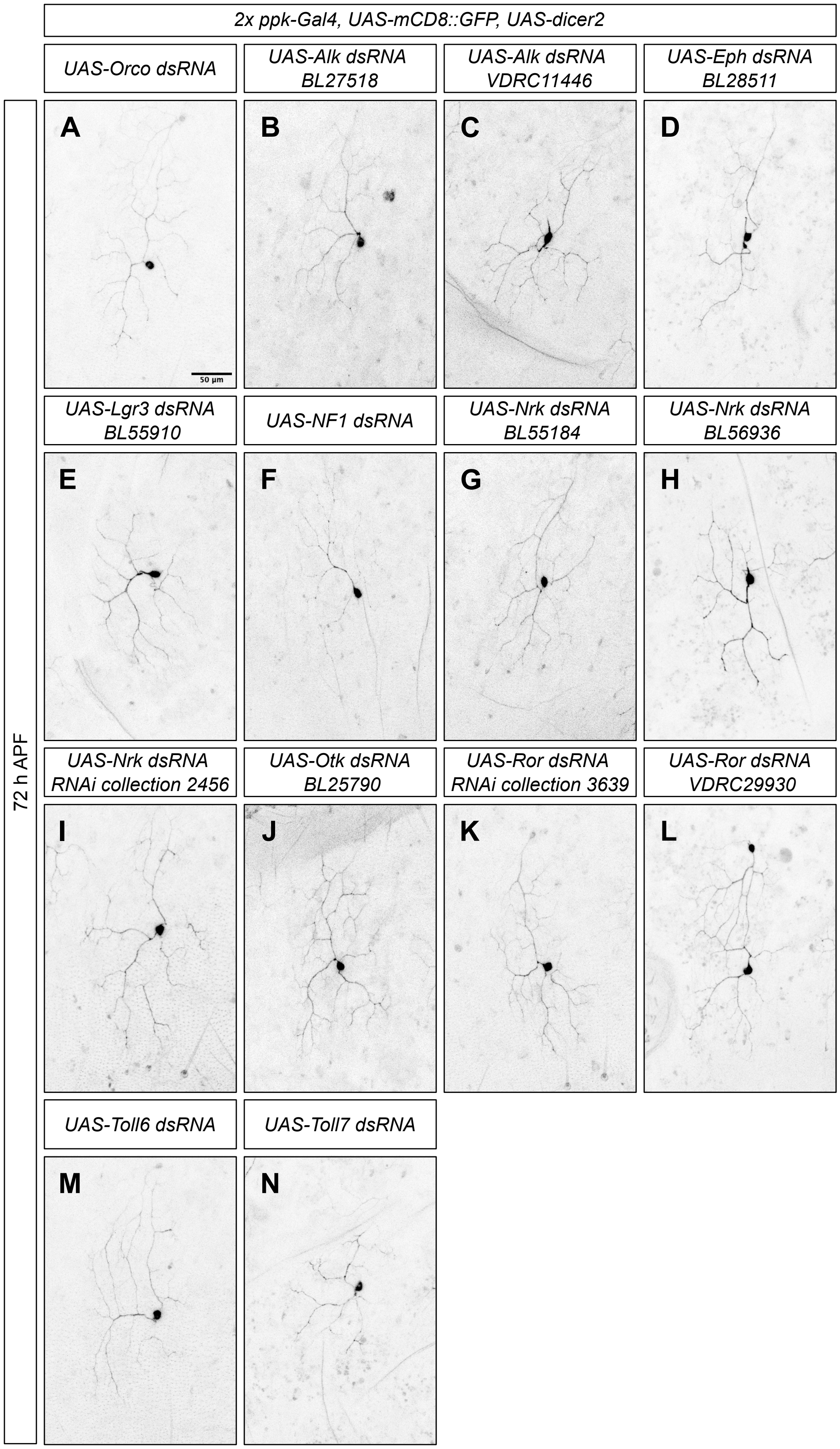

#### Sanal Figure S2

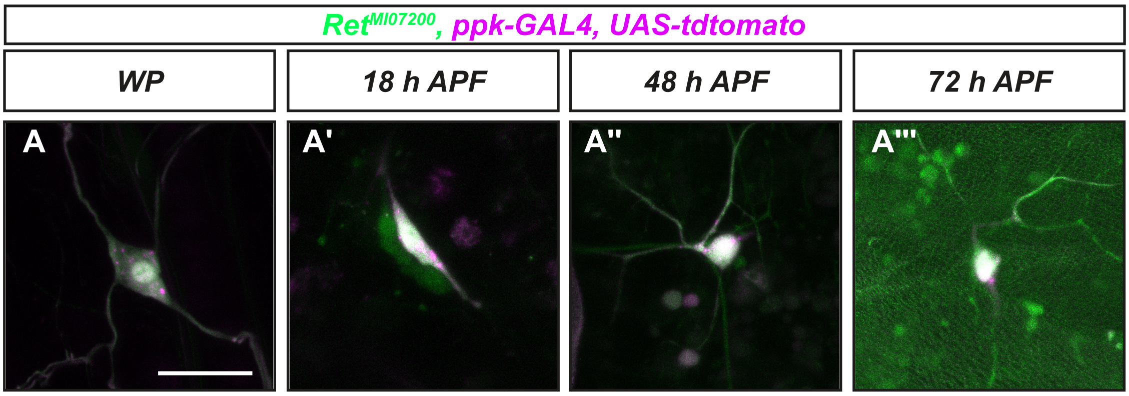

#### Sanal Figure S3

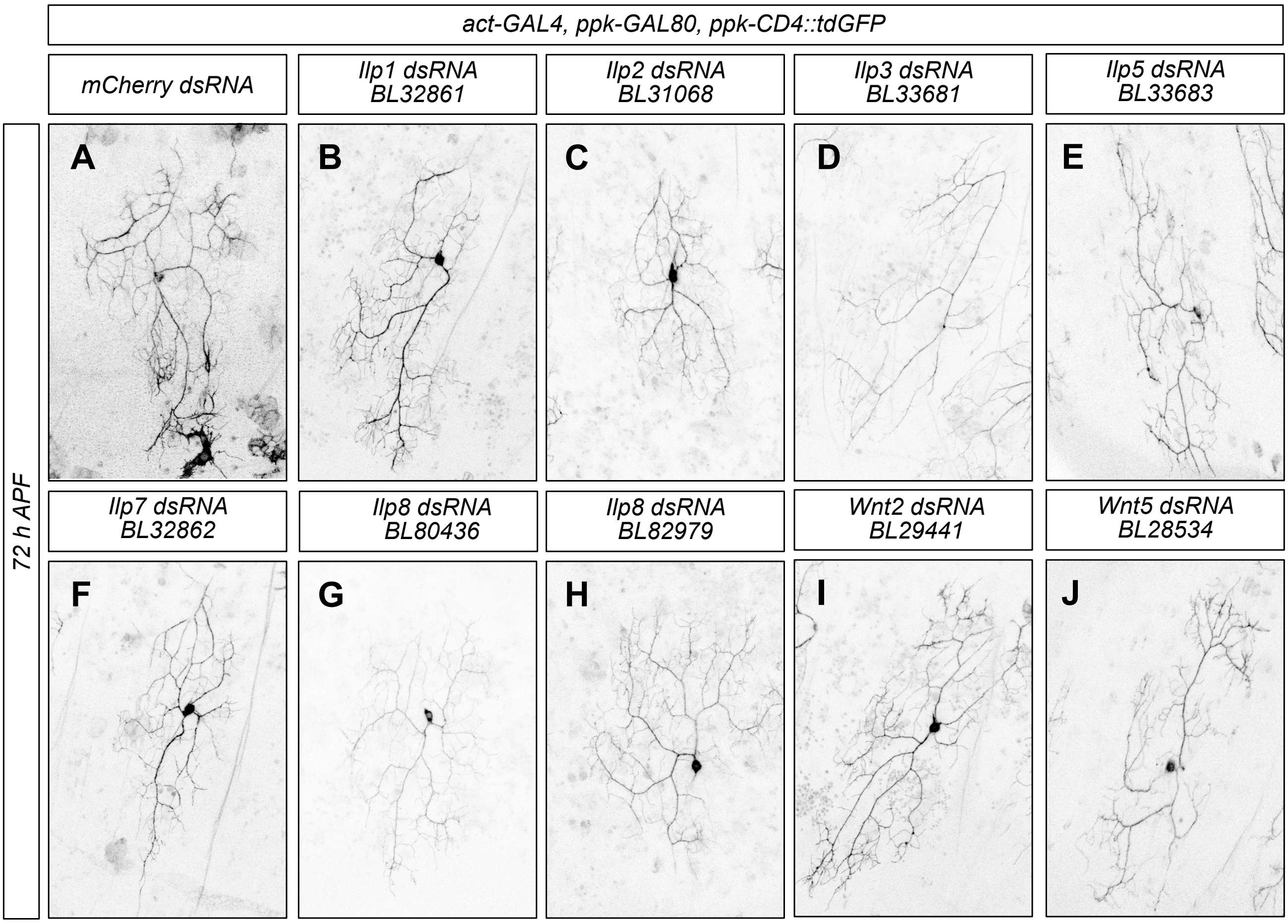

#### Sanal Figure S4

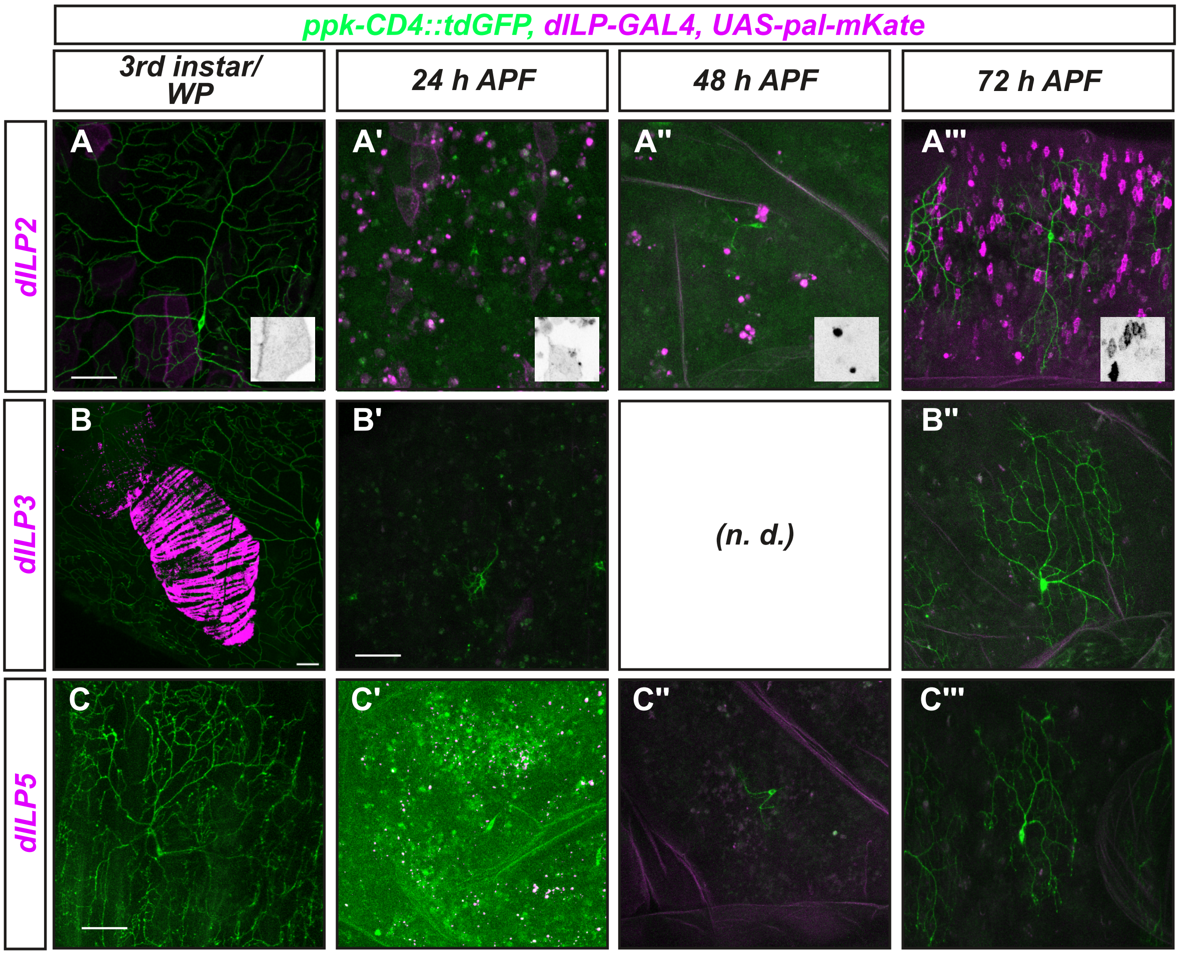
